# HA and M2 sequences alter the replication of 2013-16 H1 Live Attenuated Influenza Vaccine Infection in Human Nasal Epithelial Cell cultures

**DOI:** 10.1101/2021.11.11.468304

**Authors:** Laura M. Canaday, Jessica D. Resnick, Hsuan Liu, Harrison Powell, Alyssa M. McCoy, Dat Nguyen, Andrew Pekosz

**Affiliations:** W. Harry Feinstone Department of Molecular Microbiology and Immunology, The Johns Hopkins Bloomberg School of Public Health, Baltimore, Maryland, USA; McKusick-Nathans Institute of Genetic Medicine, Johns Hopkins University School of Medicine, Baltimore, MD 21205, USA

## Abstract

From 2013-2016, the H1N1 component of live, attenuated influenza vaccine (LAIV) performed very poorly in contrast to the inactivated influenza vaccine. We utilized a primary, differentiated human nasal epithelial (hNEC) culture system to assess the replication differences between isogenic LAIVs containing the HA segment from either A/Bolivia/559/2013 (rBol), which showed poor vaccine efficacy, and A/Slovenia/2903/2015 (rSlov), which had restored reasonable vaccine efficacy. While there were minimal differences in infectious virus production in Madin-Darby Canine Kidney (MDCK) cells, the rSlov LAIV showed markedly improved replication in hNEC cultures at both 32°C and 37°C, demonstrating that the HA segment alone could impact LAIV replication. The rSlov-infected hNEC cultures showed stronger production of interferon and proinflammatory chemokines which might also be contributing to the increased overall vaccine effectiveness of the rSlov LAIV through enhanced recruitment and activation of immune cells. The introduction of an M2-S86A mutation had no positive effects on H1 LAIV replication in hNEC cultures, in contrast to the increased infectious virus production seen with that mutation in an H3 LAIV. No obvious defects in viral RNA packaging were detected, suggesting the HA function may be driving the differential infectious virus production in hNEC cultures. The use of physiologically relevant temperatures and primary cell cultures demonstrated that candidate LAIVs can replicate efficiently, which is a necessary property for effective vaccines.

## INTRODUCTION

Live attenuated influenza vaccines (LAIV) garnered much attention when first introduced due to their potential to induce both a longer, more robust immune response and their ease of administration [1]. This is achieved using live, replicating influenza virus strains that have been attenuated to prevent illness. These viruses maintain some capacity to replicate in the upper respiratory tract to elicit a mucosal, as well as systemic, immune response. To create the influenza A/H1N1 and A/H3N2 components of the LAIV, the HA and NA gene segments from circulating virus strains are combined with the remaining internal gene segments derived from the cold-adapted master donor virus, A/Ann Arbor/6/1960 (H2N2) to generate recombinant 6:2 viruses [11]. LAIV viruses can be produced using either eggs or Madin-Darby canine kidney (MDCK) cells with similar replicative ability, although the impact of egg adaptation on vaccine efficacy has been reported [31, 21].

Virus replication after immunization elicits an immune response that more closely resembles natural infection [2]. Multiple clinical trials have shown that LAIV is more effective than the inactive influenza vaccine (IIV), even in years with antigenic drift, leading to the licensing of LAIV by the Food and Drug Administration (FDA) in 2003 for use in the United States [3].

However, the 2013-14 season LAIV vaccine showed only 13-17% effectiveness compared to 60-70% effectiveness with the antigenically matched IIV [1,8,9]. In an observational study during the 2013-14 season, 0/18 children with seronegative baseline titer against A(H1N1)pdm09 seroconverted after vaccination with LAIV compared to 7/13 who received the IIV (26). LAIV vaccine efficacy continued to decline during the 2015-16 season, where vaccine effectiveness against A(H1N1)pdm09 was not significantly different from unvaccinated children while the IIV had higher efficacy (27). The H3N2 and influenza B virus (IBV) LAIV components showed similar effectiveness when compared to IIV throughout this time [9]. Consistent effectiveness was seen for the LAIV outside the U.S. [1]. “FluMist^®^ Quadrivalent”, the LAIV used in the U.S., was manufactured by AstraZeneca and subsequently not recommended by the Advisory Committee on Immunization Practices (ACIP) before the 2016 influenza season[4,5].

A study by Hawksworth, et. al. compared the replication of successful pre-2009 LAIV strains to the less effective A(H1N1)pdm09-like strains and suggests that decreased replicative ability may lead to poor vaccine efficacy [30]. While the LAIV was not in use in the United States, studies of LAIV using a different backbone showed that by updating the H1N1 component from A/California/07/2009-like to a A/Michigan/45/2015-like strain (specifically, A/17/New York/15/5364) there was increased virus replication *in vitro*, increased virus shedding in children post-vaccination, and increased serum humoral and cellular immunogenicity, measured by seroconversion and flu-specific CD4+ T cells [28]. In 2018, the H1N1 LAIV component was officially changed, and additional efficacy testing was performed leading to an ACIP recommendation to return to using LAIV in the U.S. markets [9]. The new A/Slovenia/2903/2015 H1N1 LAIV (an A/Michigan/45/2015-like virus) displayed better thermostability and better capability to go through multiple rounds of replication in nasal epithelial cells, unlike the A/California/07/2009-like A/Bolivia/559/2013 predecessor [29, 30].

While it was originally thought that the HA and NA gene segments from any influenza A virus strain could be used with the A/Ann Arbor/6/1960 backbone to create a successful vaccine, the failure of A/California/7/2009-like virus derived vaccines suggests that the surface proteins chosen can have a significant impact on viral fitness and subsequent vaccine efficacy. The attenuation phenotype of the A/Ann Arbor/6/1960 backbone has been mapped to the PB1, PB2, and NP segments, however recent studies demonstrate that the M segment can also contribute to LAIV attenuation through an Ala to Ser mutation at M2 position 86 acquired during cold adaptation of A/Ann Arbor/6/1960; a mutation that is only found in H2N2 influenza A virus strains [12]. The A/Ann Arbor/6/1960 backbone grows optimally at 25°C and is unable to replicate above 39°C [11, 14]. It is worth noting that the upper portion of the human respiratory tract, including the nasal passageways, is at a temperature of 32°C, while the core body temperature, including the lower respiratory tract, is at 37°C [33,34,35].

We investigated the replication of H1N1 LAIVs encoding the HA segment of either the failed A/Bolivia/559/2013, the improved A/Slovenia/2903/2015, or the IIV component A/Michigan/24/2015 to assess the contribution of the H1 HA to LAIV replication *in vitro*. We demonstrate that the H1 HA from A/Bolivia/559/2013 supports less efficient LAIV replication in primary human nasal epithelial cell (hNEC) cultures when compared to A/Slovenia/2903/2015 at both 32°C and 37°C. Introducing a M2-S86A mutation, previously shown to improve H3 LAIV replication, into the H1N1 LAIVs failed to improve virus replication [12]. These data indicate that care must be taken to choose HA proteins that can promote effective LAIV replication – particularly in primary cell culture systems such as hNEC cultures - in addition to matching the HA protein antigenic structure to circulating H1N1 strains.

## MATERIALS AND METHODS

### Phylogenetic Analysis

Thirty-five HA nucleotide sequences of pdmH1N1, mainly clades 6B and 6B.1 including A/Bolivia/559/2013, A/Michigan/45/2015, and A/Slovenia/2903/2015 retrieved from GISAID (Supplementary Table) were used to build a phylogenetic tree using MEGA X software [1]. The evolutionary history was inferred by using the Maximum Likelihood method and Hasegawa-Kishino-Yano model [2]. The bootstrap consensus tree inferred from 100 replicates [3] is taken to represent the evolutionary history of the taxa analyzed [3]. Branches corresponding to partitions reproduced in less than 50% bootstrap replicates are collapsed. The percentage of replicate trees in which the associated taxa clustered together in the bootstrap test (100 replicates) are shown next to the branches [3]. Initial tree(s) for the heuristic search were obtained automatically by applying Neighbor-Join and BioNJ algorithms to a matrix of pairwise distances estimated using the Maximum Composite Likelihood (MCL) approach, and then selecting the topology with superior log likelihood value. A discrete Gamma distribution was used to model evolutionary rate differences among sites (5 categories (+G, parameter = 0.3479)). There were a total of 1701 positions in the final dataset.

### Plasmids

The plasmid pHH21, encoding each of the influenza H2N2 A/Ann Arbor/6/1960 internal gene segments, under the control of the human RNA polymerase I promoter and murine RNA polymerase I terminator [12,13], was used to generate recombinant viruses. Mutations were introduced to the M plasmid using the QuikChange Lightning site-directed mutagenesis (Agilent, Santa Clara, CA) protocol. Primer sequences are available upon request. Changes in each virus were confirmed by Sanger Sequencing. Plasmids encoding A/Slovenia and A/Bolivia HA were designed based on available sequences, cloned into pHH21, and purchased through GenScript (GenScript USA Inc., Piscataway, NJ).

### Cell Culture

Madin-Darby canine kidney (MDCK) and human embryonic kidney 293T (293T) cells were cultured in Dulbecco’s Modified Eagle Medium (DMEM, Sigma-Aldrich) with 10% fetal bovine serum (FBS, Gibco Life Technologies), 100U penicillin/mL with 100μg streptomycin/mL (Quality Biological), and 2mM L-Glutamine (Gibco Life Technologies) at 37°C with air supplemented with 5% CO_2_. Infectious medium (IM) was used in all infections and consists of DMEM with 4ug/mL N-acetyl trypsin (NAT), 100u/ml penicillin with 100ug/ml streptomycin, 2mM L-Glutamine and 0.5% bovine serum albumin (BSA) (Sigma). Human nasal epithelial cell (hNEC) cultures were isolated from non-diseased tissue after endoscopic sinus surgery for non-infection related conditions [12]. The cells were collected from 4 female patients. The cells were differentiated at an air-liquid interface (ALI) in 24-well Falcon filter inserts (0.4-μM pore; 0.33cm^2^; Becton Dickinson) before infection, using ALI medium on basolateral side of insert, as previously described [14].

### Recombinant Viruses

Recombinant viruses were rescued using a 12 plasmid reverse genetics system [12, 13]. 293T cells were transfected with 0.5μg of pHH21 plasmids encoding H2N2 A/Ann Arbor/6/1960 LAIV internal genes PB2, PB1, PA, NP, NS, and the WT or M2-S86A mutant M. 0.5μg of A/Michigan/45/2015, A/Bolivia/55/2013, or A/Slovenia/2903/2015 HA and 0.5μg of A/Michigan/45/2015 NA in the pHH21 plasmid were added to supply the surface proteins. Additionally, 1μg of protein expression plasmids for A/Udorn/72 PB2, PB1, and NP plus 0.2μg PA were added as plasmids that would reconstitute the influenza polymerase activity. After a 24-hour incubation at 37°C with 5% CO_2_, 5×10^5^ MDCK cells in 100μL of IM were added. Cells were sampled daily until obvious signs of cytopathic effects were visible, at which time the supernatants were harvested, cell debris pelleted at 500g for 5 minutes, and the clarified supernatant stored at −80°C.

### Plaque Assay

To generate virus stocks, 90-100% confluent 6-well plates of MDCK cells were inoculated with 250μL of serial 10-fold dilutions of transfection supernatant in IM and incubated at 32°C with 5% CO_2_ for one hour, with gentle distribution of the solution every 15 minutes. Wells were then covered with 1% agarose in 1X MEM plus 1:1000 NAT. Once the agarose solidified, plates were incubated at 32°C with 5% CO_2_ for 5 days. Plaques were picked using a 1mL large bore pipette tip, added to tubes containing 0.5 ml IM, and stored at −80°C to be used to establish seed stocks of that monoclonal virus colony (see below). To determine plaque morphology, the above protocol was performed using sequence verified virus stocks. After 5 days of incubation, plates were fixed with 4% formaldehyde in PBS overnight and stained with Naphthol Blue Black overnight. To quantify plaque area, images of the wells were captured using a dissecting microscope with an Olympus DP-70 color camera and plaque area measurements were captured via ImageJ (NIH).

### TCID_50_ Assay

MDCK cells were grown to 90-100% confluence in 96-well plates. After being washed twice with PBS+ (Phosphate Buffered Saline containing 0.9 mM Calcium and 0.5 mM Magnesium), ten-fold serial dilutions of the viruses in IM were made, and 20μL of each dilution was added to 6 wells. The plates were incubated at 32°C with 5% CO_2_ for 7 days. The cells were fixed by adding 100μL of 4% formaldehyde in PBS per well for at least 1 hour, supernatant discarded, and then stained with Naphthol Blue Black solution overnight. Endpoint values were calculated by the Reed-Muench method [15].

### Seed Stocks and Working Stocks

To generate seed stocks, fully confluent 6-well plates of MDCK cells were washed 2x with PBS+ and then infected with 250μL of the plaque pick solution for 1 hour, with redistribution every 15 minutes, at 32°C with 5% CO_2_. After 1 hour, the infection media was removed, and 2mL of IM was added. Virus supernatant was collected when 75% of the cells showed cytopathic effect, typically 5 days post-infection. The seed stocks were titrated via TCID_50_ assay. Working stocks were generated from seed stocks, in a similar fashion, except on fully confluent MDCK cells in 75cm^2^ flasks, and the seed stock inoculum was diluted to a multiplicity of infection (MOI) of 0.001 TCID_50_s per cell in IM.

### Low Multiplicity of Infection (MOI) Growth Curve (GC)

An MOI of 0.01 TCID_50_ per cell was used in MDCK cells. The virus inoculum was diluted in IM, 100μL was added to the cells in 24 well tissue culture plates in triplicate and allowed to incubate at 32°C or 37°C for 1 hour, with redistribution every 15 minutes. The inoculum was removed, the cells were washed 3x with PBS+, and incubated with 500μL of IM. At the indicated times, all media was collected, stored at −80°C, and fresh IM was added.

For hNEC cell infections, an MOI of 0.1 TCID_50_s per cell was used. The basolateral media was collected, stored at −80°C, and replaced with fresh media. The apical side of the trans-well was washed 3 times with IM (without NAT), with a 10-minute 32°C or 37°C incubation between each wash. The virus inoculum was diluted in IM, 100μL was added to the cells, and allowed to incubate at 32°C or 37°C for 2 hours. The inoculum was removed, the cells washed 3 times with PBS without Mg^+2^ or Ca^+2^ and incubated at 32°C or 37°C with 5% CO_2_. At the indicated times, IM was added to the apical side, allowed to incubate at the corresponding temperature for 10 minutes, and then collected and stored at −80°C. Basolateral media was collected, stored at −80°C, and replaced every 48 hours. Infectious virus particle production in apical washes was quantified using TCID_50_ on MDCK cells.

### Cytokine Expression

Secreted interferons, cytokines, and chemokines were quantified from the basolateral samples from 48- and 96-hours post infection from the hNEC multi-step virus growth curves. Measurements were performed using the V-PLEX Human Chemokine Panel 1 (CCL2, CCL3, CCL4, CCL11, CCL17, CCL22, CCL26, CXCL10, and IL-8) (Meso Scale Discovery) and the DIY Human IFN Lambda 1/2/3 (IL-29/28A/28B) ELISA (PBL Assay Science) according to the manufacturer’s instructions. Each sample was analyzed in duplicate, and expression was normalized to mock infected cultures from the same donor.

### Competition Assay

hNEC wells were acclimated to 32°C for 4 hours prior to being co-infected with the indicated virus strains. To match the 0.1 MOI used in growth curves, the TCID_50_ units of rSlov WT working stock need for 0.05 MOI was converted to RNA/μL, and all other viruses were normalized to that value (actual resulting MOIs ranged from 0.1-0.5 due to different RNA: TCID_50_ ratios between viruses). Infections were performed as described for growth curve assay above. RNA was extracted from 70 μL of apical wash combined with 70 μL of PBS using the Qiagen vRNA extraction mini kit and converted to cDNA using a Universal Flu A primer as previously described [32]. RNA segment copies/ μL was then determined using digital droplet PCR (ddPCR). The same forward and reverse primers were used for each set, and the probes were designed to target differences in either M2 or HA coding region as described, with the mutated residue in the middle (Table 1). All primers and probes were ordered from BioRad. 5 μL of cDNA was used for each reaction regardless of concentration. Droplet generation was performed according to manufacturer protocol with BioRad Qx200 Droplet Generator using droplet generation oil for probes. PCR cycling conditions were as follows: 95°C for 10min, 40 cycles of 94°C for 30s and 53°C for 1min, 98°C for 10min. Droplet reading was performed according to manufacturer protocol with BioRad QX200 Droplet Reader. Percentage of each indicated segment was then determined based on number of FAM or HEX positive droplets divided by the total number of positive droplets measured for each sample.

**Table 1.**
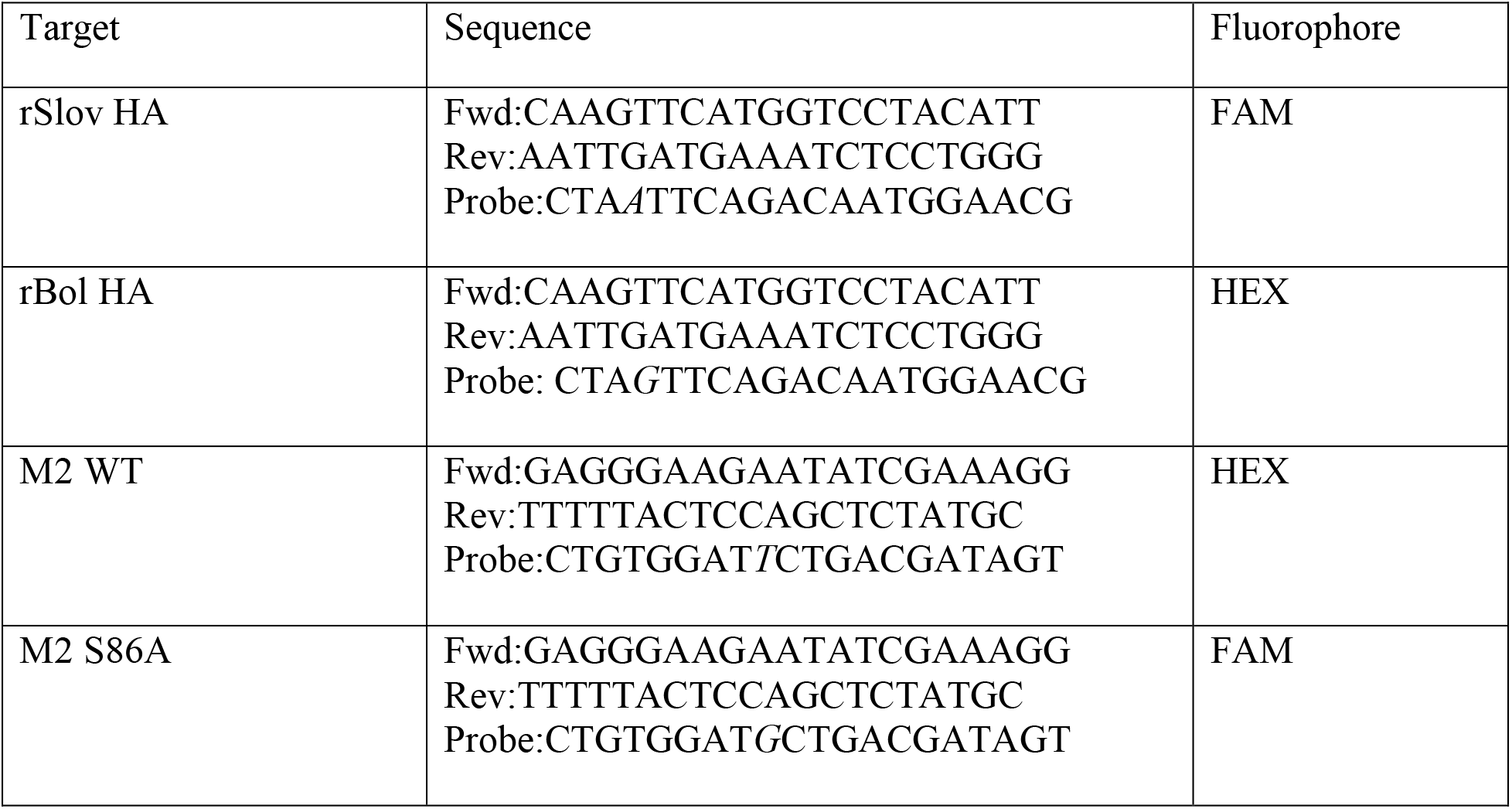
ddPCR primers utilized.

## RESULTS

### Rescue of recombinant H1 LAIVs and replication in MDCK cells

An alignment of the A/California, A/Bolivia, A/Slovenia, and A/Michigan HA protein sequences was performed to identify all amino acid changes (Fig. 1A). When compared to A/California, a majority (11/16) of the changes were seen across all 3 HA proteins. Q223R is a mutation detected in egg-adapted A/Michigan. When the A/Slovenia and A/Bolivia proteins are directly compared, only 4 amino acid differences are identified (Fig. 1A, pink, italicized amino acids). The mutation at position 162 shows the gain of a conserved glycosylation site in A/Slovenia and A/Michigan viruses, and residue 216 is an amino acid that aids in the recognition of sialic acid in the HA receptor binding site [16,17]. Phylogenetic analysis (Fig. 1B) shows that A/Bolivia and A/Slovenia represent the 6B and 6B.1 clades respectively. The 6B.1 clade is defined by amino acid changes S84N, S162N and I216T, which account for 3 of the 4 differences between the two vaccine viruses [18]. Recombinant LAIVs encoding the HA proteins from A/Bolivia (rBol), A/Slovenia (rSlov), and A/Michigan (rMich) were generated using plasmid based infectious clone technology.

**Fig. 1.**
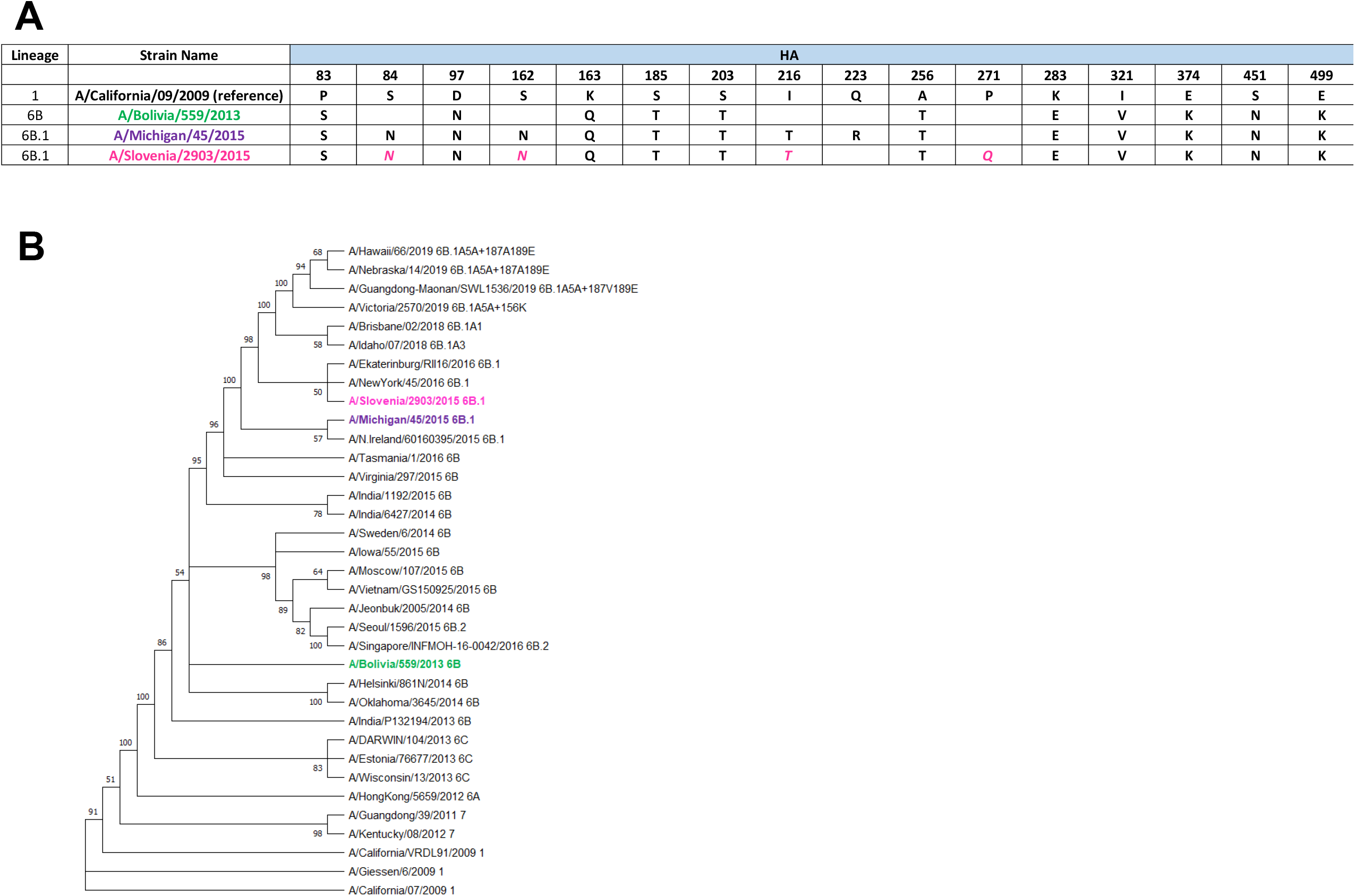
Alignment and phylogenetic tree of HA sequences from the H1N1 LAIV viruses used. (A) Alignment schematic of H1 vaccine virus HA protein sequence. Amino acid differences compared to the A/Cal reference strain are shown. Amino acid differences between A/Bolivia and A/ Slovenia are shown in pink italics. Phylogenetic tree (B) of IAV pdmH1N1 HA gene sequences. The tree was constructed by using the maximum-likelihood method and the HKY+G model in MEGA X software. Numbers to the left of nodes are bootstrap values (100 replicates). Tips are strain names and followed by its clade. HA sequences of thirty-five H1N1 viruses were retrieved from GISAID. The three viruses used in the study are in bold and colored.

At both 32°C, a temperature consistent with the upper respiratory tract in humans, and 37°C, the temperature of the lower portion of the human respiratory tract, rSlov had faster kinetics of infectious virus production in MDCK cells, though all three viruses reached the same peak infectious virus titer (Fig. 2A and 2B). Plaque assays performed at 32°C and 37°C on MDCK cells also show no statistically significant difference between the three viruses at the same temperature, or with any one virus compared across temperatures (Fig. 2C). Together, these data indicate that the three viruses show comparable, high amounts of infectious virus production and replication in MDCK cells at both temperatures, with rSlov having a somewhat faster replication rate.

**Fig. 2.**
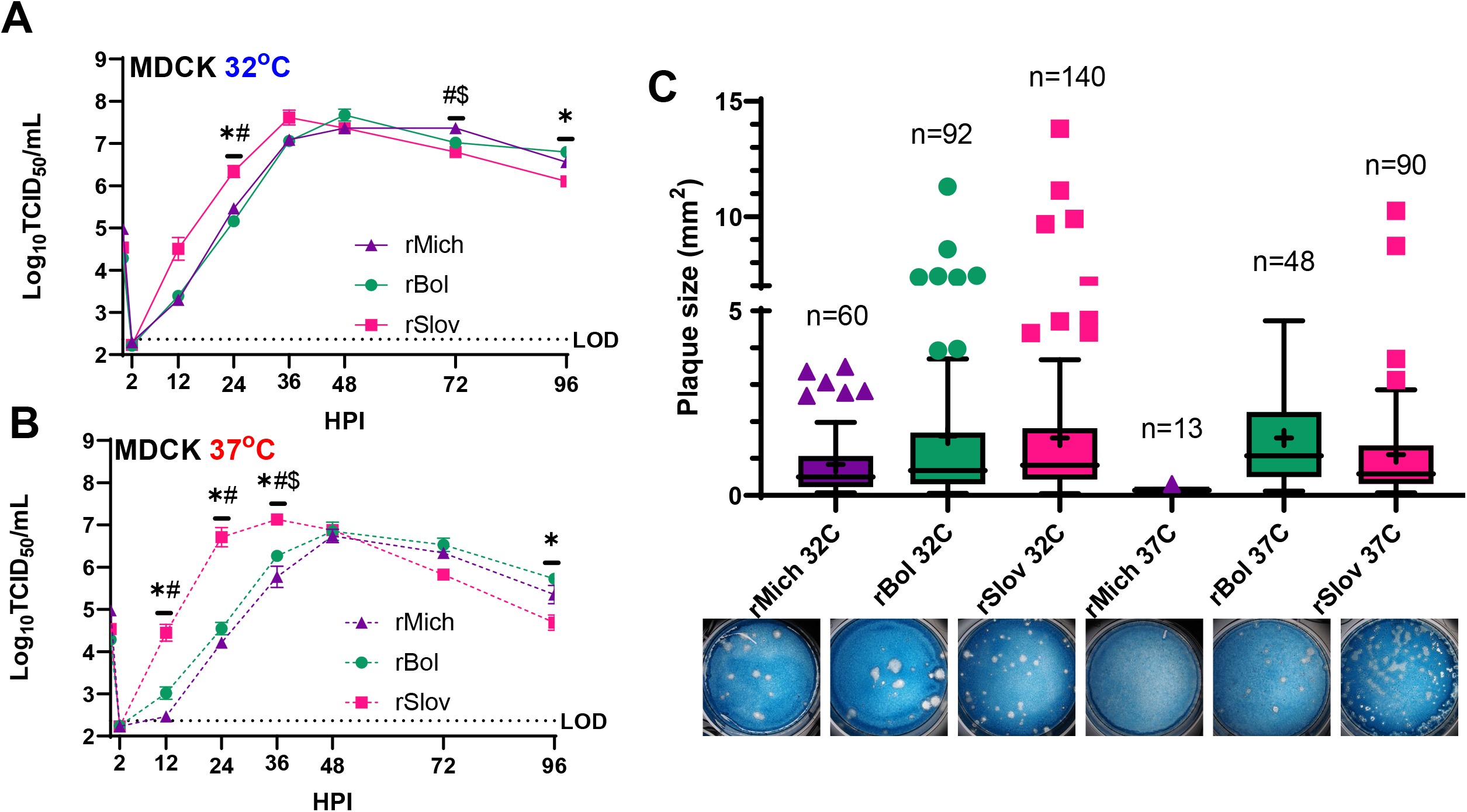
Replication of H1N1 LAIVs on MDCK cells at different temperatures. Low MOI, multistep growth curves were performed on MDCK cells at 32oC (A) and 37oC (B) with the indicated viruses. Data are pooled from three independent replicate experiments each with n=3 wells per virus (total n = 9 wells per virus, SEM). Significant differences are indicated as follows: *rSlov v rBol, #rSlov v rMich, $ rBol v rMich, p < 0.05 (two-way repeated measures ANOVA with Tukey’s posttest, only 12-48hr included). Dotted line indicates limit of detection. (C) Plaque assays were performed with the indicated viruses on MDCK cells. Plaque diameter was quantified from 20-140 individual plaques per virus identified from 3 independent experiments at 32o or 37oC. *p < 0.05 (one-way ANOVA with Tukey’s posttest).

### Replication of WT viruses on hNEC cultures

We utilized human nasal epithelial cell (hNEC) cultures collected from non-diseased patients in the Johns Hopkins Medical Institutions as a second in vitro system to assess LAIV replication, as they have previously been shown to be more sensitive to changes in LAIV replication compared to MDCK cells [12,14,23]. Low MOI growth curves were performed on these cultures at both 32°C and 37°C. At 32°C, rSlov showed faster replication kinetics and replicated to a higher peak titer than rBol or rMich (Fig. 3A) – with a striking 100- to 1000-fold difference in infectious virus production compared to rBol. The rBol virus had slower initial replication kinetics compared to rMich, but it reached a similar peak titer. At 37°C, rSlov showed the greatest amount of infectious virus production, with peak infectious virus production approximately 1000-fold below what was seen at 32°C, consistent with phenotypes noted previously [12,14]. At this temperature, rBol failed to produce any detectable infectious virus, suggesting a severe, temperature-dependent reduction in virus fitness (Fig. 3B). Together these data indicate that rSlov has the greatest capacity to replicate on hNECs at both 32°C and 37°C, compared to rMich and rBol.

**Fig. 3.**
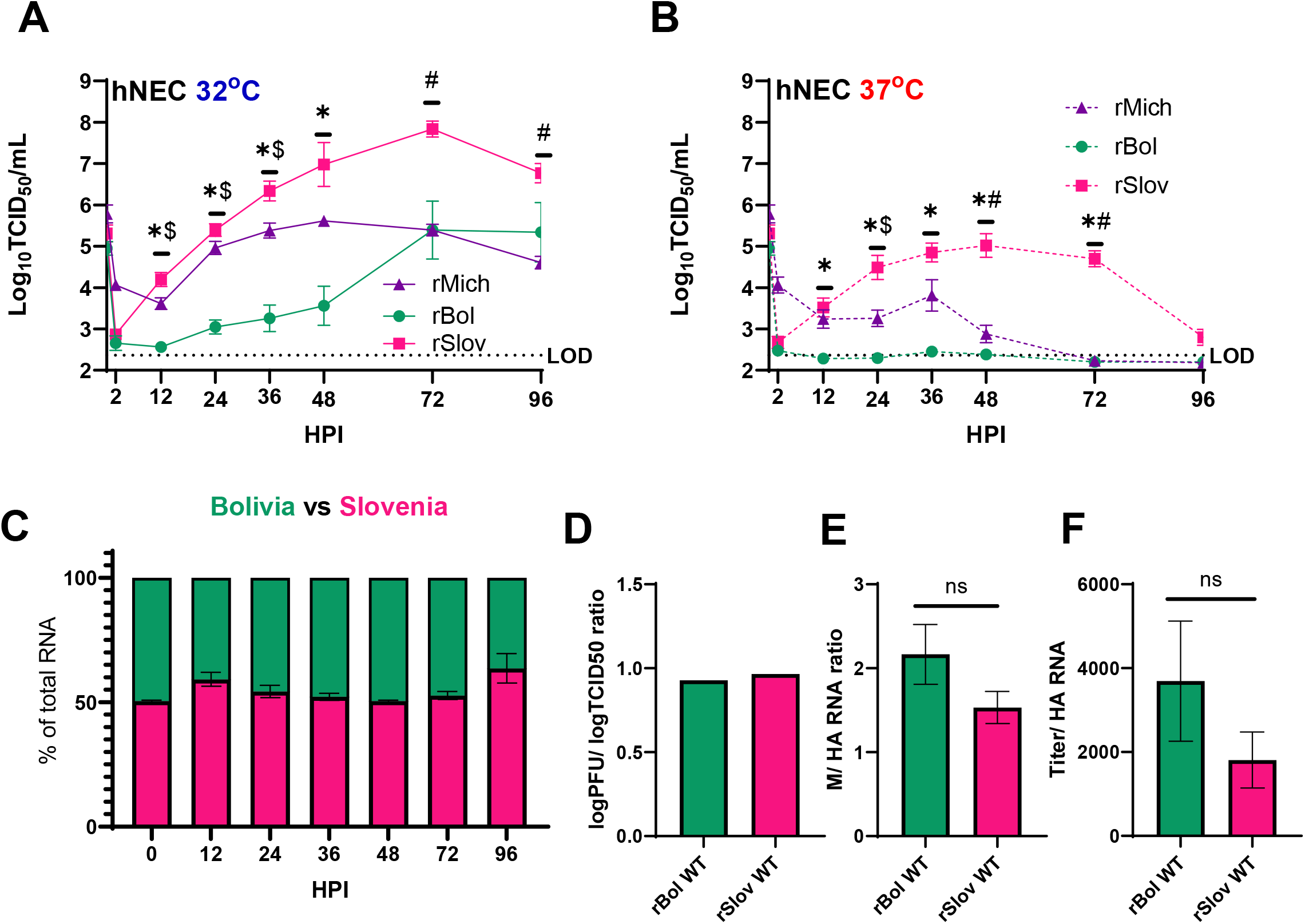
Replication of H1N1 LAIVs on hNECs. Low MOI multistep growth curves were performed on hNECs at 32oC (A) and 37oC (B) with the indicated viruses. Data are pooled from three independent replicate experiments each with n = 4 wells per virus (total n = 12 wells per virus). *rSlov v rBol, #rSlov v rMich, $ rBol v rMich, p < 0.05 (two-way ANOVA with Tukey’s posttest, SEM). Dotted line indicates limit of detection. Competition growth assays were performed with the indicated viruses on hNECs at 33°C and percent of total vRNA was determined by ddPCR (C). Data are pooled from three independent experiments each with n=3 wells (9 wells total, SEM). (D) Ratio of determined TCID50/mL to PFU/mL for each indicated virus from MDCK grown virus stocks. Virus titer was performed in duplicate and PFU/mL was determined from 6 wells of previously shown plaque assays. M to HA segment RNA ratio was also calculated (E) using ddPCR on 13 10-fold virus dilutions. These same dilutions were then used to calculate the TCID50/mL to HA RNA ratio (F) for each virus to predict differences in infectious vs non-infectious virus production (SEM). P< 0.05(one-way ANOVA with Tukey’s posttest.

To directly compare the fitness of viruses encoding the rBol and rSlov HA segments, we developed a digital droplet PCR (ddPCR) assay that could differentiate the rBol and rSlov HA segments, then used that assay to assess HA segment packaging in hNEC cultures co-infected with rBol and rSlov. We infected hNEC cultures with equal amounts of rBol and rSlov normalized to the input RNA. Over the course of the growth curve, no difference in the amount of rSlov or rBol HA RNA was detected, suggesting that these two RNA segments are packaged at the same frequency in newly produced virus particles (Fig. 3C).

Using the MDCK grown viruses, we confirmed that there was no difference in detection of infectious virus using either the TCID_50_/mL or PFU/mL methods (Fig. 3D), then compared the amounts of HA and M segment RNA and found no significant differences in the ratio of packaged RNA (Fig. 3E). Finally, we compared the ratio of HA segment RNA concentration to TCID_50_/mL and found no significant differences (Fig. 3F), suggesting the viruses produced from MDCK cells were not different in their inherent infectivity or RNA packaging.

### Replication of M2-86 Mutants on MDCK cells at 32°C and 37°C

Changing M2-86 from a Serine residue to an Alanine residue in and LAIV encoding the H3N2 A/Victoria/361/2011 (rVic) virus HA and NA segments increased LAIV infectious virus production in hNEC cultures but not MDCK cells, suggesting this mutation contributes to the LAIV attenuation phenotype [12]. We hypothesized that introducing this mutation into the rBol virus would increase virus replication to levels consistent with rSlov in hNEC cultures. Recombinant viruses encoding M2-S86A with all three H1 HA proteins were generated and characterized for replication in MDCK cells. The replication of rVic M2-WT and rVic M2-S86A was not significantly different at 32°C, but the S86A virus produced slightly more infectious virus at 37°C (Fig. 4A and 4B), consistent with previous observations [12]. In contrast, M2-S86A had no effect on rMich infectious virus production at either temperature (Fig. 4C and 4D), slightly improved rBol replication at 32°C (Fig. 4E and 4F), and decreased rSlov infectious virus production at both temperatures (Fig. 4G and 4H). These data suggest the M2-S86A mutation has limited, but strain specific, effects on H1N1 viruses when grown on MDCK cells [12].

**Fig. 4.**
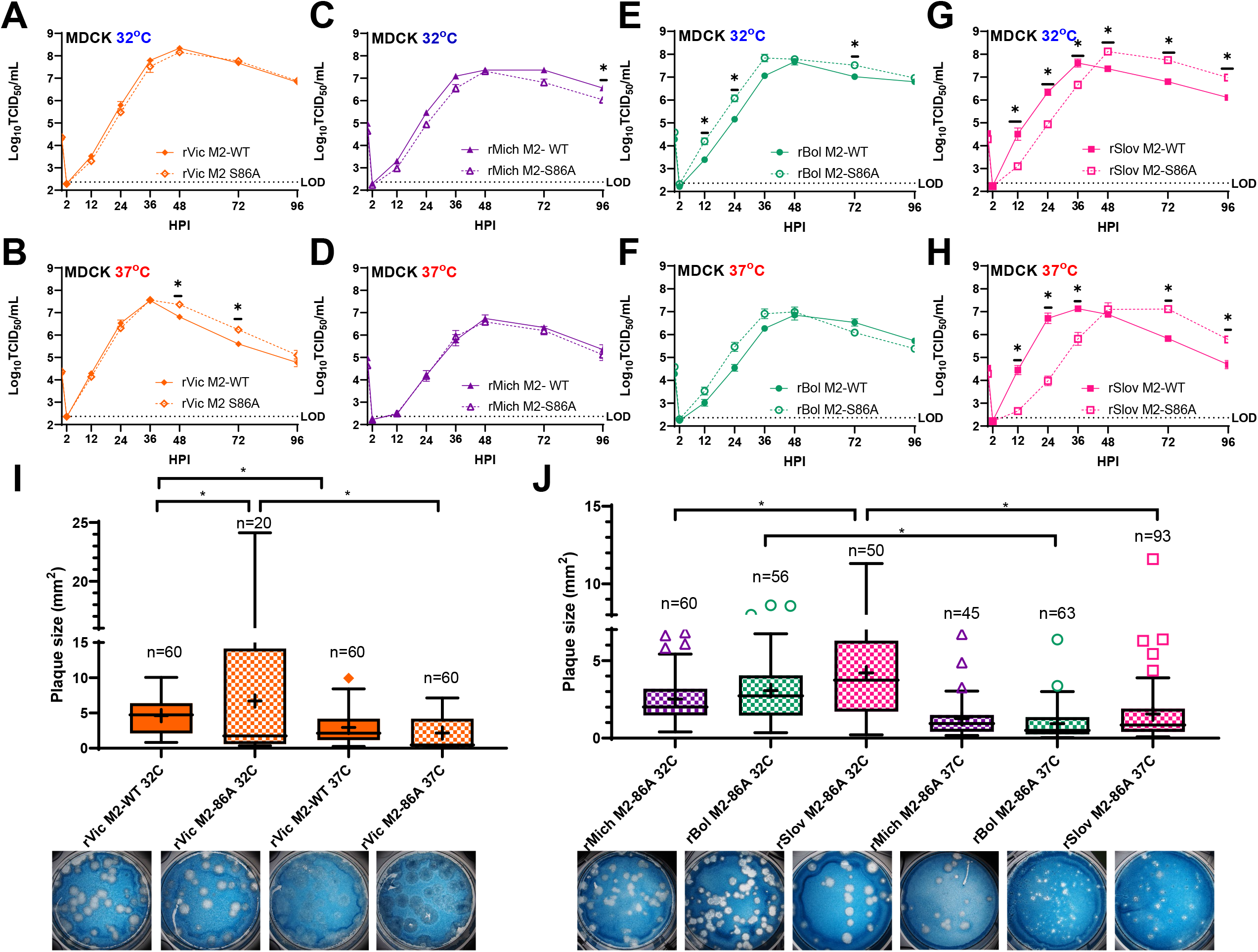
M2 mutations that enhance H3N2 LAIV replication have variable affects on H1N1 LAIV replication in MDCK cells. Low MOI, multistep growth curves were performed at 32oC (A,C,E,G) and 37oC (B,D,F,H) with the indicated viruses. Data are pooled from three independent replicate experiments each with n=3 wells per virus (total n = 9 wells per virus, SEM). * p< 0.05 (two-way repeated measures ANOVA with Tukey’s posttest, only 12-48hr included). Dotted line indicates limit of detection. (I) and (J) Plaque assays were performed with the indicated viruses on MDCK cells. Plaque diameter was quantified from 24-93 individual plaques per virus identified from 3 independent experiments at 32oC or 37oC. *p < 0.05. (one-way ANOVA with Tukey’s posttest).

Plaque assays of rVic showed a significant increase in plaque size for rVic M2-S86A at 32°C compared to 37°C and M2-WT 32°C (Fig. 4I). rVic M2-WT generated larger plaques at 32°C than at 37°C. rSlov M2-S86A at 32°C generated the largest plaques of all 3 H1 HA viruses across both temperatures (Fig. 4J). Notably, all 3 H1 viruses generated larger plaques at 32°C than at 37°C. All 3 H1 viruses with M2-S86A generated larger plaques than with M2-WT at 32°C (Fig. 2C). These data indicate that the M2-S86A mutation may improve cell-to-cell spread in a temperature dependent manner, but the M2-S86A mutation does not show a temperature dependent effect on infectious virus production on MDCK cells.

### Replication of M2-S86A Mutants on hNEC cultures at 32°C and 37°C

The M2-S86A mutation had its greatest effect on H3N2 LAIV replication in hNEC cultures [12]. In low MOI growth curves using rVic M2-WT and M2-S86A on hNECs, no replication differences were seen at 32°C, while a statistically significant increase in infectious virus production was seen at 37°C (Fig. 5A and 5B), consistent with previous reports [12]. For the rMich viruses, the M2-86A mutation resulted in decreased infectious virus production at both 32°C and 37°C, with nearly no infectious virus detected at 37°C with the M2-S86A mutation (Fig. 5C and 5D). The rBol LAIV M2-WT and M2-S86A had no differences in virus production at either temperature, with no infectious virus production at 37°C (Fig. 5E and 5F). In the rSlov LAIV, the M2-S86A mutation resulted in reduced infectious virus production at both temperatures, with no infectious virus production seen at 37°C (Fig. 5G and 5H). Since all three H1 LAIVs rescued with the M2-S86A mutation were unable to replicate at 37°C, the data indicate an HA subtype/strain and temperature specific role for this mutation in LAIV replication in hNEC cultures.

**Fig. 5.**
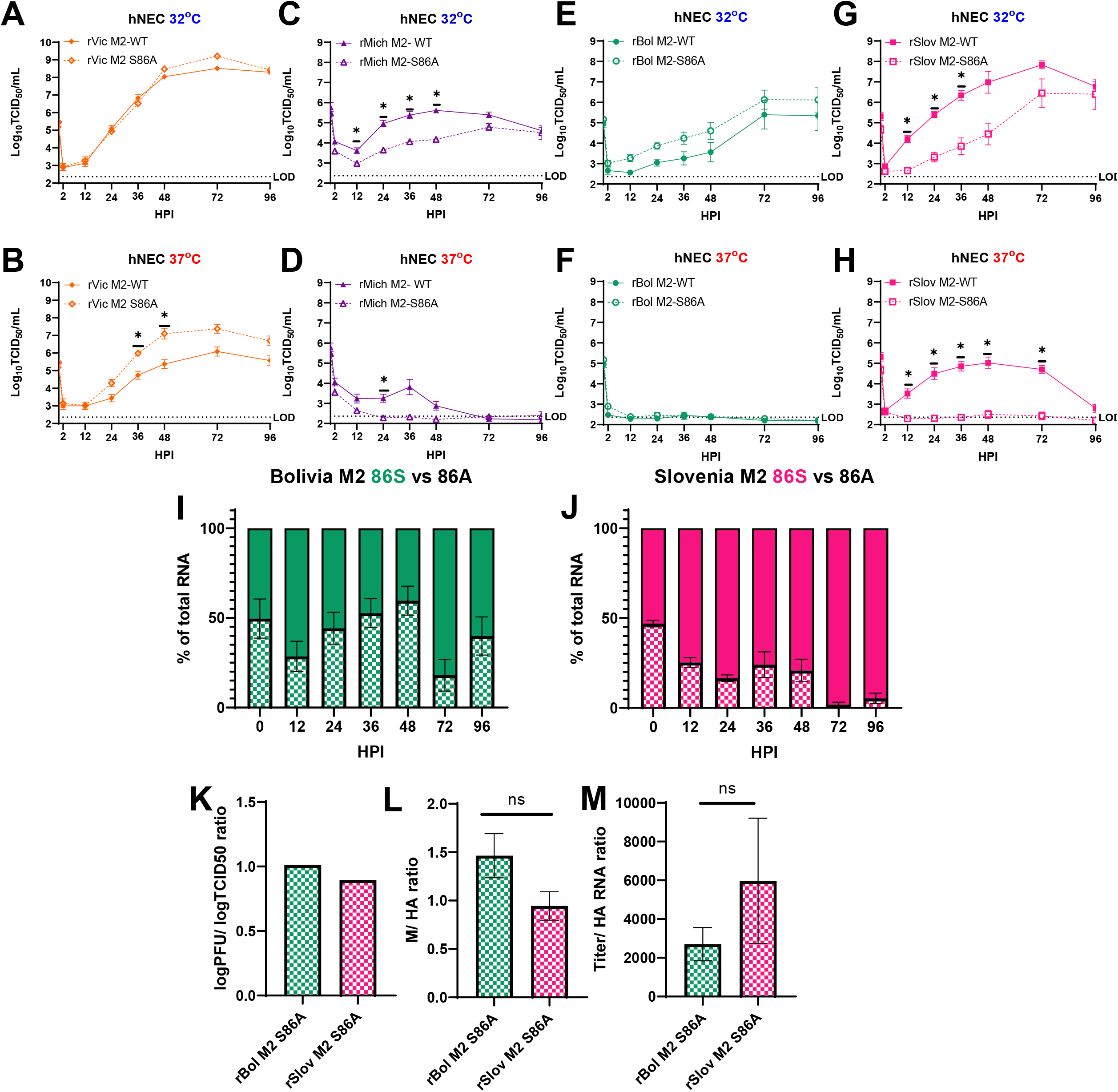
M2 mutations that enhance H3N2 LAIV replication reduce H1N1 LAIV Replication in hNEC cultures. Low MOI, multistep growth curves were performed at 32oC (A,C,E,G) and 37oC (B,D,F,H) with the indicated viruses. Data are pooled from three independent replicate experiments each with n=4 wells per virus (total n = 12 wells per virus, SEM). *p < 0.05 (two-way repeated measures ANOVA with Tukey’s posttest). Dotted line indicates limit of detection. Competition growth assays were performed with the indicated viruses on hNECs at 33°C and percent of total vRNA was determined by ddPCR (I,J). Data are pooled from three independent experiments each with n=3 wells (9 wells total, SEM). (K) Ratio of determined TCID50/mL to PFU/mL for each indicated virus from MDCK grown stock viruses. Infectious virus titer was performed in duplicate and PFU/mL was averaged from 6 wells of previously shown plaque assays. M to HA segment RNA ratio was also calculated (L) using ddPCR on 13 10-fold virus dilutions. (M) These same dilutions were then used to calculate the TCID50/mL to HA RNA ratio for each virus to predict differences in infectious vs non-infectious virus production (+SEM). P< 0.05(one-way ANOVA with Tukey’s posttest).

We performed a ddPCR competition assay in hNEC cultures at 32°C for both the rSlov and rBol viruses encoding either M2-WT or M2-S86A. in agreement with the infectious virus production in the growth curve (Fig. 5E), roughly equal amounts of M2-WT and M2-S86A RNA in rBol co-infected cultures were detected over the course of the experiment (Fig. 5I). For rSlov, we again see that M2-WT has an advantage (Fig. 5J), consistent with what was observed in the infectious virus growth curve (Fig. 5G). For the MDCK-grown stocks of the rBol M2-S86A and rSlov M2-S86A, there were no differences in PFU/mL to TCID_50_/mL ratios (Fig. 5K), and no statistically significant differences in either M segment to HA ratio or the ratio of infectious virus to HA RNA segment (Fig. 5L, Fig. 5M). Overall, we conclude that the M2-S86A mutation does not broadly improve virus replication for H1N1 LAIV viruses as it did for H3N2 [12].

### rSlov virus elicits a stronger innate immune response than rBol in hNEC cultures

Basolateral secretion of interferons, cytokines, and chemokines involved in the inflammatory response to viral infection were measured at both 48- (Fig. 6A) and 96- (Fig. 6B) hours post-infection (hpi) with WT viruses at 32°C. At 48 hpi, rSlov elicited a larger total innate immune response, with significantly higher expression of both CXCL10 and CCL11 (Fig. 6A). This was stronger at 96 hpi, with rSlov eliciting significantly higher expression (over 10-fold) of IFN-lambda, CCL17, CCL13, CCL2, CXCL10, CXCL8, and CCL11 (Fig. 6B). In contrast, expression of 8/11 cytokines and chemokines measured were significantly higher in rSlov-WT infected samples, there was significantly higher induction of CCL3 in rBol-WT infected cultures at 96 hpi. Overall, this data suggests that level of innate immune mediator induction was different between the two H1N1 LAIV viruses and could contribute to differential immune responses.

**Fig. 6.**
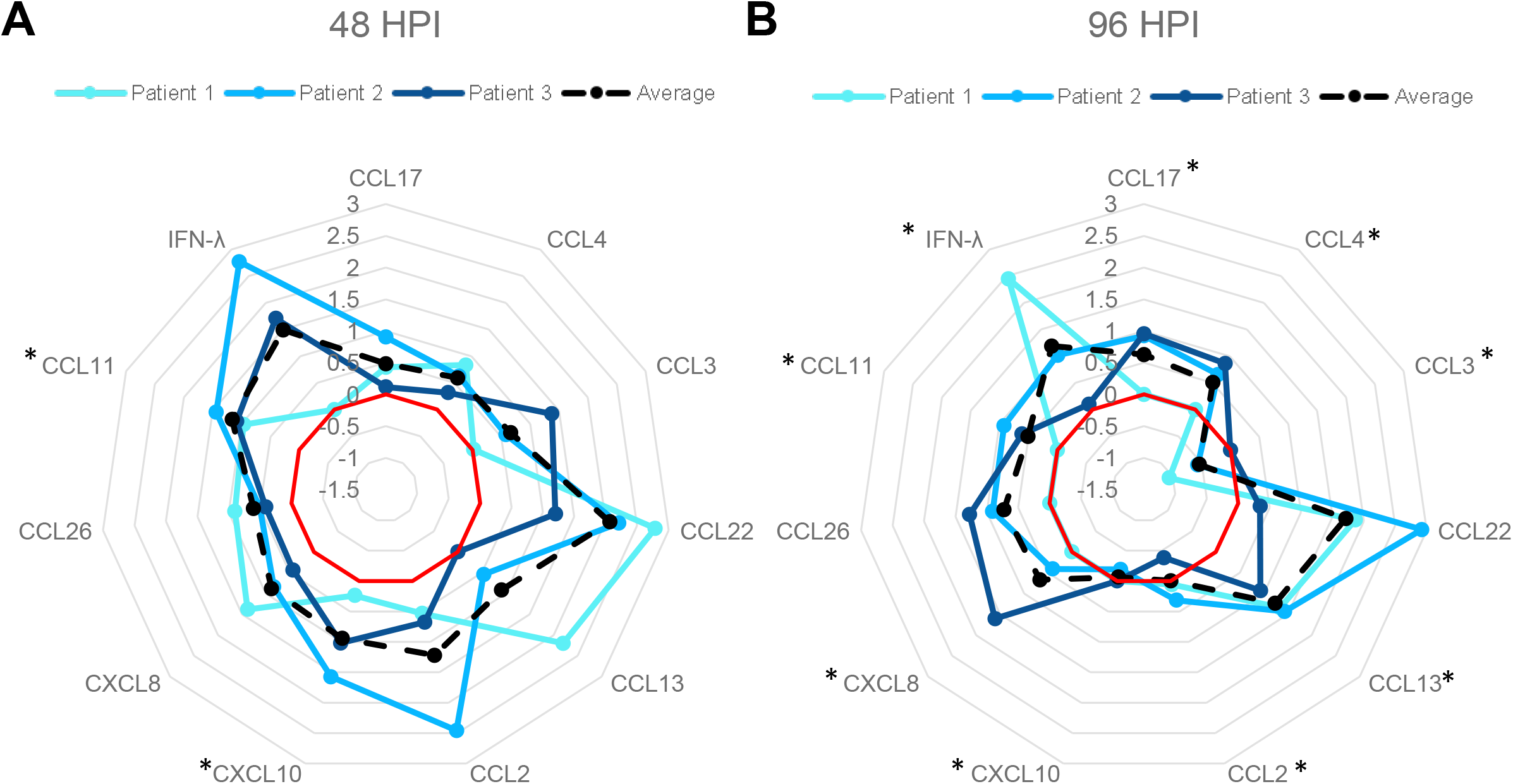
Epithelial cell innate immune responses after infection with rSlov or rBol in hNEC cultures. Basolateral secretion of interferons, cytokines and chemokines was measured at 48 (A) and 96 (B) HPI during low MOI multistep growth curve experiments on hNECs at 32oC. Log fold change of rSlov relative to rBol is shown for each individual hNEC culture and as an average of all experiments. The red line indicates log fold change of 0 or no difference. Values outside red circle indicate higher expression in rSlov infected wells while values inside the red circle indicate higher expression in rBol infected wells. *p < 0.05. (two-way repeated measures ANOVA with Tukey’s posttest).

## DISCUSSION

A/California/7/2009-like LAIV vaccines showed lower vaccine efficacy than both their antigenically matched IIV or later A/Michigan/24/2015-like LAIVs [26,27,28,29,30]. The decrease in vaccine efficacy is hypothesized to be due to an inability of the virus to undergo the multiple rounds of replication in the upper respiratory tract required to induce a strong immune response [28,29,30]. This is most apparent with the 2015-2016 season LAIV donor A/Bolivia/559/2013 which was shown to have decreased replication in immortalized cell lines compared to the improved 2017-2018 season LAIV donor virus A/Slovenia/2903/2015 [9,19].

In this study, we show that while successful LAIV viruses, such as A/Slovenia, show subtle fitness differences on immortalized cell lines, they replicate significantly better on the physiologically relevant primary human nasal epithelial cell cultures and show less severe temperature sensitivity. These data are consistent with what was seen by AstraZeneca, the manufacturer of FluMist, when they conducted these experiments after rBol failed as a vaccine [9]. We showed the utility of primary cell cultures to assess LAIV virus replication [12,14], and other groups also demonstrated their utility for evaluating LAIV replication [20]. However, our observations of the effect of physiological ranges of temperature as an additional factor impacting LAIV replication in hNEC cultures and the contribution of HA segments to the temperature sensitivity phenotype show the strength of using hNEC cultures at physiological temperatures as a tool to assess LAIV replication fitness. Previous work showed that changes in the HA protein resulting from the embryonated egg adaptation process that vaccine viruses go through can affect both the HA antigenic structure and overall fitness of the resulting virus [21], so perhaps it is not surprising that as few as a four amino acid difference in the HA segments can cause such dramatic differences in LAIV replication.

Additionally, we show for the first time HA subtype specific mutations in the A/Ann Arbor/ 6/1960 LAIV backbone that influence temperature sensitivity in hNEC cultures. Previous work showed that an M2-S86A mutation increases H3N2 viral replication at higher temperatures on hNECs [12]. However, here we show that that the same mutation either has no effect or decreases H1 LAIV viral fitness. Our data suggests that the differences in fitness observed between both the M2-WT and M2-S86A viruses may be due to differences in infectious particle production. Mutations at the M2 86 residue contribute to defective interfering particle production [22], but our analysis of viral RNA packaging did not suggest such an effect was taking place in these H1 LAIVs. Additionally, LAIVs overall have been shown to produce large amounts of defective-interfering viral RNAs as a way to reduce infectivity, but this alone does not drive the reduction in vaccine effectiveness [24, 25, 36]. Taken together, our data suggests that the HA segment chosen for H1N1 LAIVs is an important determinant of innate immune induction, viral replication and perhaps subsequent vaccine efficacy.

Testing candidate LAIVs for replication on hNEC cultures and over a range of physiological temperatures allows for the detection of differences in viral replication that may correlate with vaccine efficacy [23]. The additional power of the hNEC system is the ability to measure innate immune induction in response to infection, which we show follows viral replication in magnitude and is greater overall in more successful vaccine viruses. We propose that hNEC cultures should be included in future LAIV screening pipelines, and that special consideration should be given to the HA segment chosen for future vaccines, as even small amino acid changes can have drastic effects on replication. Future work should identify H1N1 subtype specific mutations to improve the LAIV backbone and allow for more diverse HA options.

## ACKNOWLEDGEMENTS

We thank the members of the Andrew Pekosz, Sabra Klein, and Kimberly Davis laboratories for insightful comments and discussions pertaining to this manuscript. The work was supported by the National Institute of Allergy and Infectious Diseases grants HHSN272201400007C (Johns Hopkins Center of Excellence in Influenza Research and Surveillance, AP), N7593021C00045 (Johns Hopkins Center of Excellence in Influenza Research and Response, AP), 5T32GM007814-37 (JR), and T32 AI007417 (HP, JR and AMM). Other support was provided by the Eliasberg Family Foundation. We thank the Bloomberg Flow Cytometry and Immunology Core for use of ddPCR and MSD instruments.

